# Functional ultrasound (fUS) imaging of displacement-guided focused ultrasound (FUS) neuromodulation in mice

**DOI:** 10.1101/2024.03.29.587355

**Authors:** Seongyeon Kim, Nancy Kwon, Md Murad Hossain, Jonas Bendig, Elisa E. Konofagou

## Abstract

Focused ultrasound (FUS) stimulation is a promising neuromodulation technique with the merits of non-invasiveness, high spatial resolution, and deep penetration depth. However, simultaneous imaging of FUS-induced brain tissue displacement and the subsequent effect of FUS stimulation on brain hemodynamics has proven challenging thus far. In addition, earlier studies lack in situ confirmation of targeting except for the magnetic resonance imaging-guided FUS system-based studies. The purpose of this study is 1) to introduce a fully ultrasonic approach to in situ target, modulate neuronal activity, and monitor the resultant neuromodulation effect by respectively leveraging displacement imaging, FUS, and functional ultrasound (fUS) imaging, and 2) to investigate FUS-evoked cerebral blood volume (CBV) response and the relationship between CBV and displacement. We performed displacement imaging on craniotomized mice to confirm the in targeting for neuromodulation site. We recorded hemodynamic responses evoked by FUS and fUS revealed an ipsilateral CBV increase that peaks at 4 s post-FUS. We saw a stronger hemodynamic activation in the subcortical region than cortical, showing good agreement with the brain elasticity map that can also be obtained using a similar methodology. We observed dose-dependent CBV response with peak CBV, activated area, and correlation coefficient increasing with ultrasonic dose. Furthermore, by mapping displacement and hemodynamic activation, we found that displacement colocalizes and linearly correlates with CBV increase. The findings presented herein demonstrated that FUS evokes ipsilateral hemodynamic activation in cortical and subcortical depths and the evoked hemodynamic responses colocalized and correlate with FUS-induced displacement. We anticipate that our findings will help consolidate accurate targeting as well as an understanding of how FUS displaces brain tissue and affects cerebral hemodynamics.

## Introduction

Focused ultrasound (FUS) can modulate excitatory and inhibitory neurons [1,2] and has been extensively investigated in the central (CNS) [1-15] and peripheral nervous system (PNS) [16-18] with advantageous characteristics of high spatial resolution and deep penetration depth [19]. The biophysical and cellular mechanisms of FUS neuromodulation in CNS have been elucidated at a length, that is known via mechanosensitive ion channel [20-22]. FUS generates the acoustic radiation force that exerts on neuronal tissue, induces microscopic displacement, and eventually activates the channel. Thus, understanding how FUS displaces structures in the brain is important to shed light on mechanical mechanism of FUS and to ensure successful neuromodulation.

One of the major challenges in developing FUS in preclinical and clinical applications is in situ confirmation of FUS targeting. Earlier preclinical studies mostly relied on the assumption of beam propagation in the free field [6,9,11,13] or simulation to account for skull aberration [12,14]. However, this approach is not ideal, given that the brain is a viscoelastic and inhomogeneous structure with viscoelasticity even modulating according to neural activity [23]. Without in situ confirmation of targeting, FUS is likely bound to be subject to errors in alignment or positioning of the FUS transducer, which eventually may result in a discrepancy in neuromodulation outcome [5,19]. Mohammadjavadi et al. (2022) [15] leveraged magnetic resonance acoustic radiation force imaging (MR-ARFI) to non-invasively measure microscopic displacement induced by FUS in sheep with intact skulls. They also found that displacement can be used to evaluate neuromodulation efficacy affected by different skull attenuation between subjects. However, the technique requires a large, expensive, complex, and not portable system, thus making it difficult for researchers to easily adopt the system into their FUS experimentation. Therefore, we sought to develop a system more compatible with the FUS setup for in situ targeting. We previously developed and demonstrated displacement imaging for peripheral nerve stimulation [16-17]. Here in this study, we leveraged displacement imaging for in situ targeting in the mouse brain during FUS.

Cerebral hemodynamics grants a readout of brain activity via neurovascular coupling. The investigation of brain hemodynamics in response to FUS is important to validate, characterize, and optimize the neuromodulatory effect of FUS on CNS. Multiple earlier studies have demonstrated that FUS can causally modulate brain activity leading to motor responses [6,9,11-14,25,36]. Yet, only a few studies have elucidated the effect of FUS on cerebral hemodynamics [7,26,27]. Yuan et al. (2020) [26] demonstrated cortical hemodynamic response induced by FUS and its dependency on FUS parameters, but the measurement was confined to the cortex and lacks a hemodynamic map which is important to shed light on neuromodulatory impact of FUS at the network level and changes in functional connectivity in the brain [28].

Functional ultrasound (fUS) imaging can measure hemodynamics with high spatiotemporal resolution [29]. Being analogous to functional MRI (fMRI), fUS utilizes the changes in cerebral blood volume (CBV) as a correlate of neural activity [30]. In sharp contrast with optical methods, fUS is capable of imaging the whole brain in 2D or 3D and, in contrast to fMRI, can operate in real time, measuring more transient signals with high spatiotemporal resolution. Recently, we demonstrated cortical hemodynamic responses in healthy and neuropathic mice in response to peripheral FUS stimulation [24]. We herein switched our interest into CNS to elucidate hemodynamics in response to central FUS stimulation.

Here, we introduce the fully ultrasonic approach to in situ target, modulate neural activity and monitor FUS-mediated neuromodulation by leveraging displacement imaging, FUS, and fUS imaging, respectively. We demonstrated displacement imaging of the brain in craniotomized mice, FUS-evoked cerebral activity from cortical to subcortical depth, and its dose-dependency on ultrasonic parameters. By utilizing high frequency (4 MHz) FUS, we were able to demonstrate more targeted neuromodulation with the lateralization of FUS-evoked hemodynamic responses via fUS imaging. Lastly, we first map displacement and hemodynamic response to provide displacement-activation map and elucidate the colocalization and correlation between displacement and CBV.

## Materials and methods

### 2.1. Animal preparation

All experiments and procedures in this study were performed in accordance with Columbia University Institutional Animal Care and Use Committee (Protocol # AC-AABF2550; approval on 2023-03-30). Female C57BL/6J (Envigo; Indianapolis, IN, USA) ages 8 – 12 weeks were used in all experiments. In the study, a total of six wild-type mice (n = 4: fUS imaging with FUS, n = 1: histological evaluation, n = 1: temperature measurement) were used. Animals were anesthetized using isoflurane (3% for induction; 2 – 2.5% during preparation, craniotomy; 0.8 - 1% during displacement imaging and fUS imaging with FUS). In all animals, a large-window (9 mm by 5 mm) craniotomy was conducted for higher signal-to-noise ratio (SNR) in displacement and fUS imaging. Body temperature was monitored and kept at 37°C with a heating pad with a rectal temperature sensor. Animals were subjected to toe pinch every 15 min to assess the depth of anesthesia and showed a modest response to toe pinch at 0.8% - 1% isoflurane. Isoflurane % was adjusted to achieve normal breathing without gasping.

### 2.2. Experimental setup

Fig. 1A depicts the experimental setup used in this study to perform displacement imaging and fUS imaging with FUS neuromodulation. We used a 128-element linear imaging transducer (L22-14vXLF; Vermon, France) for displacement and fUS imaging. The transducer was connected to an ultrasound research system (Vantage 256 High-Frequency Option; Verasonics Inc., Kirkland, WA, USA). Matlab (Mathworks, Natick, Massachusetts, USA) was used to program custom transmit and receive acquisitions. With regard to FUS stimulation, Verasonics ultrasound system sends a trigger to a function generator (33550B Waveform Generator, Keysight, Santa Rosa, CA, USA) in a synchronized manner. The function generator amplified by a RF amplifier (A150; E&I, Rochester, NY, USA) drove a FUS transducer. The single-element 4 MHz FUS transducer (H-215; SonicConcepts, Bothell, WA, USA) was coaxially aligned with the imaging transducer through a 3D-printed attachment and was used to generate FUS for displacement and neuromodulation. The transducers are positioned using a 3D-motorized positioning system (BiSlide; Velmex, Bloomfield, NY, USA). A 3D-printed collimator with an acoustic-permeable membrane was filled with degassed water to couple the transducers with the craniotomized brain.

**Figure 1.**
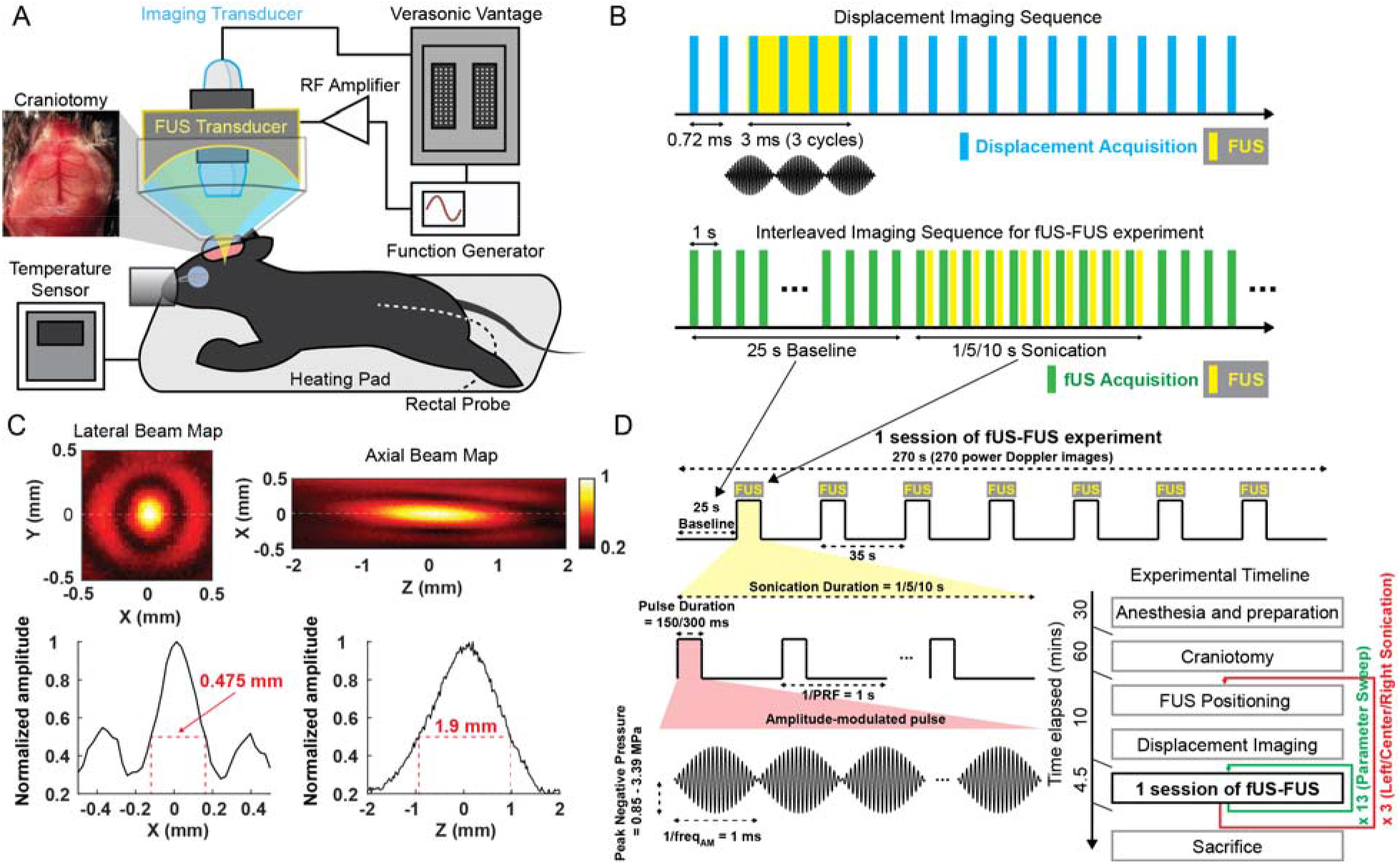
(A) Experimental setup for displacement imaging and functional ultrasound (fUS) imaging with FUS neuromodulation. (B) Displacement imaging sequence and interleaved imaging sequence for fUS-FUS experiment. (C) Hydrophone measurement of the axial and lateral beam profile of the 4 MHz FUS transducer (top) and corresponding full-width at half-maximum (bottom). (D) fUS-FUS experiment and FUS stimulation protocol (top and left, respectively), and experimental timeline (right bottom).

### 2.3. Preparation of large-window craniotomy

A rectangular cranial window (9 mm by 5 mm) was made about 10 minutes before a first session of the fUS-FUS experiment started. The head of the animal was positioned in a stereotaxic system (SGM-4; Narishige, Tokyo, Japan), and the hair of the head was removed using a clipper and depilatory cream. A single incision was made and skin was removed to expose the skull so that the skull sutures were visible. A thin wire was placed at Bregma 0 mm and imaged with B-mode to zero the imaging transducer relative to Bregma coordinates. After removing connective tissue on the surface of the skull, a micromotor drill with a foot pedal (51449; Stoelting Co, Wood Dale, IL, USA) was used to carefully drill out the window centered around Bregma. Synthetic interstitial fluid (SIF) [31] was applied on the skull frequently to cool down the skull and prevent bleeding. When the skull window is ready to be lifted, it is carefully removed by fine forceps. Once the brain was exposed, SIF-soaked gauze was placed on the exposed brain. The SIF-soaked gauze was not removed until applying ultrasound coupling gel and the transducers on the brain for experimentation.

### 2.4. Imaging plane and FUS targeting

Displacement and fUS imaging with FUS were performed at Bregma -0.5 mm (AP: -0.5 mm). As stated previously, aligning transducers to the plane of Bregma -0.5 mm was possible by zeroing the imaging transducer to the Bregma coordinate. Once aligned, 3D-rendered brain vasculature allowed fine adjustments to be made to select the imaging plane. To demonstrate the spatial resolution of FUS neuromodulation with 4 MHz FUS used in this study, three targets were explored by positioning the transducers laterally; left (ML: -1.5 mm), center (ML: 0 mm), and right (ML: 1.5 mm) sonication with the same depth of FUS focus (DV: 3 mm) at Bregma - 0.5 mm. Target engagement was assessed by displacement imaging before fUS imaging and the transducers were repositioned with the 3D positioning system till displacement was present at the intended target.

### 2.5. Displacement imaging

Displacement imaging was used to confirm successful delivery of FUS at the target and to estimate FUS-induced brain tissue displacement. The setup and method established in our previous work on a peripheral nerve [16] was modified to better estimate displacement in the brain and are reported in Supplementary Table S1. Displacement was acquired and processed with GPU-accelerated delay-and-sum (DAS) beamformer. The real-time displacement imaging enabled us to achieve in situ confirmation of targeting using displacement imaging before conducting subsequent fUS-FUS experiment.

RF data for each displacement frame was collected every 0.72 ms seconds and a 3ms FUS pulse was triggered before 3rd frame acquisition to induce displacement such that displacement trace spans for 1.42 ms before, 2.88 ms during, 8.64 ms after FUS (Fig. 1B). After computing 1D normalized cross-correlation on compounded and beamformed RF, interframe displacement was obtained. Cumulative displacement was generated by accumulating interframe displacement over the course of time. The pixel-wise displacement was averaged within full-width half-maximum (FWHM) to generate the averaged displacement. Note that, except for Fig. 2C, interframe displacement was used in all analyses, figures and videos reported in the study.

**Figure 2.**
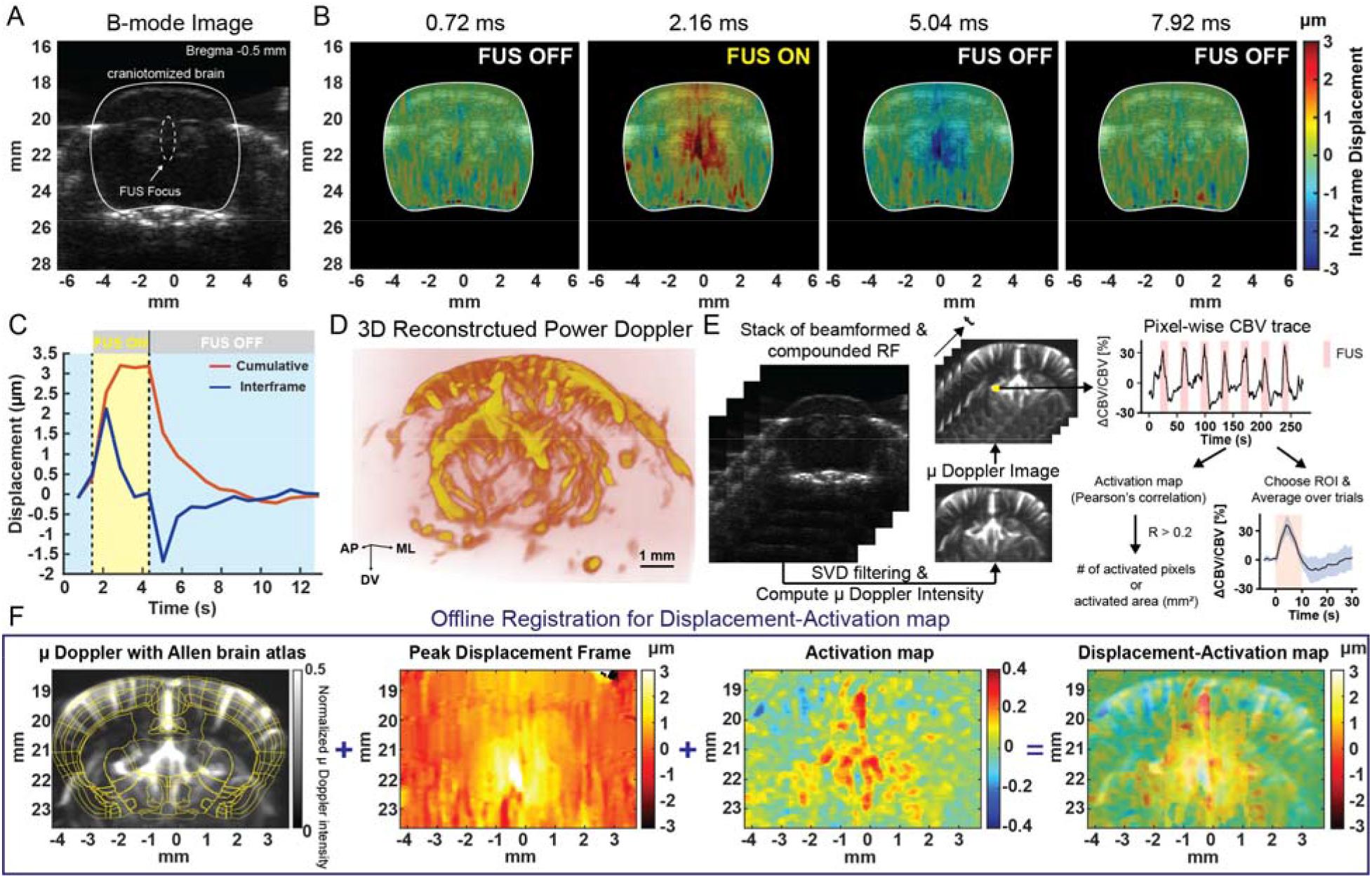
(A) B-mode image of craniotomized mouse brain at bregma -0.5 mm (coronal). FUS beam is located around the midline (center sonication). (B) Displacement imaging of the brain. The push and relaxation from FUS sonication were captured with displacement imaging, indicating successful imaging of acoustic radiation force and displacement engagement of the brain tissue. (C) Interframe (blue) and cumulative (red) displacement trace averaged within full-width half-maximum (FWHM) of displacement. (D) 3D reconstruction of the power Doppler image. (E) Processing steps to generate power Doppler images and to analyze CBV responses evoked by FUS. (F) Offline registration to create displacement-activation map.

### 2.6. FUS neuromodulation paradigms

After displacement targeting, fUS imaging with FUS neuromodulation was performed. During a session of the fUS-FUS experiment, mice were sonicated 7 times with 25 ∼ 34 s interval using the 4 MHz FUS (Fig. 1D top and left). Three FUS parameters (PNP: Peak negative pressure, SD: Sonication duration, PD: Pulse duration) were explored over 13 sessions and this set of sessions was repeated per target (Fig. 1D right bottom). PNP ranges from 0.85 to 3.39 MPa while other parameters were fixed (SD = 10 s, PD = 300 ms). In the same manner, SD ranges from 1 to 10 s (PNP = 3.39 MPa, PD = 300 ms) and PD ranges from 150 to 300 ms (PNP = 3.39 MPa, SD = 10 ms). The combination of ultrasonic parameters according to the session is reported in Supplementary Table S2. The FWHM of FUS focal size is 0.475 mm x 1.9 mm (Fig. 1C). Regarding pulsing scheme, we decided to use amplitude-modulated regime at 1 kHz amplitude modulation (AM) frequency to minimize auditory confounding that could occur during FUS sonication [3]. Note that, all peak negative pressure values reported in this study was derated to account for attenuation in brain tissue using the following equation [32]:

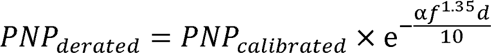

where α is 0.75 dB*cm*^-1^*MHz*^-1^ through the brain tissue [33] (assuming the linear dependence of attenuation on the frequency), f is the carrier frequency of the FUS in MHz (f = 4 MHz), and d is the depth of FUS focus in the brain in mm (d = 3 mm).

### 2.7. fUS imaging

Ultrasound imaging sequence consists of 39 tilted (3 cycles) plane waves. 3 plane waves per angle were transmitted at 13 angles evenly spaced between ± 7° and RF from the 3 planes waves was averaged internally in the ultrasound research system. Plane waves were sent at 19500 Hz (500 Hz compounded PRF). Each received RF stack consists of 150 compound frames. After acquiring and GPU DAS-beamforming compounded RF stacks, a spatiotemporal filter using the singular value decomposition (SVD) [34] was used to remove stationary tissue signals and reconstruct blood signals. Components with an eigenvalue below 30 were discarded. The imaging parameter is reported in Supplementary Table S1. Lastly, the reconstructed stack of beamformed RF was summed to obtain the final CBV image at 1 Hz functional framerate. 3D volumetric power Doppler image was acquired by 1D raster scan in 0.1 mm steps (Supplementary Video S1.2) and was rendered using open source ParaView (Kitware, Clifton Park, New York, USA) (Fig. 2D, Supplementary Video S1.1).

To prevent introducing any artifact arising from FUS interference into the fUS imaging, an interleaved sequence for FUS trigger and fUS imaging was used and controlled by the ultrasound research system (Fig. 1B). FUS was triggered once fUS acquisition ended so that GPU processing was completed during FUS. This interleaved scheme successfully prevented introducing FUS interference into power Doppler images.

### 2.8. Functional image processing

Fig. 2E depicts the overall process to obtain the activation map and CBV response using functional images. Sequences of power Doppler, or CBV images were acquired: 25 frames for baseline, 1 – 10 frames during the stimuli (dependent on sonication duration), and 34 – 25 frames for after cessation of FUS stimulation. ΔCBV/CBV was calculated pixel-by-pixel using the following equation:

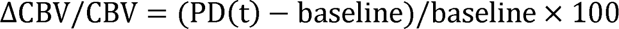

where PD is a power Doppler signal moving-averaged (window size: 4 frames) and baseline is an average of PD in all the baseline frames. Activation map was constructed pixel-by-pixel using Pearson’s correlation coefficient between CBV signal and the binary stimuli signal. Correlation values above r > 0.2 (z > 3.3 and p < 0.005 with 270 time points) were chosen as statistically significant. Final correlation map was acquired after median filtering (window size: 0.245 mm x 0.245 mm). To compute mean CBV response, two ROIs (window size: 0.343 mm x 0.343 mm) centered to the highest correlation pixel in cortex and subcortex were chosen (Fig. 3D). Mean CBV response was obtained by averaging ΔCBV/CBV of all pixels in the ROIs over 7 trials. The size of activated area was computed by multiplying the number of the activation pixels by the size of power Doppler pixel (0.0024 *mm*^2^). Lastly, displacement was registered onto activation map with the background image of power Doppler to obtain displacement-activation map (Fig. 2F, Supplementary Video S2). Note that, for registration purpose, displacement was depicted with hot color map and only displacement greater than 0.75 µm was displayed in all displacement-activation maps, but all pixels were used in the statistical analysis.

**Figure 3.**
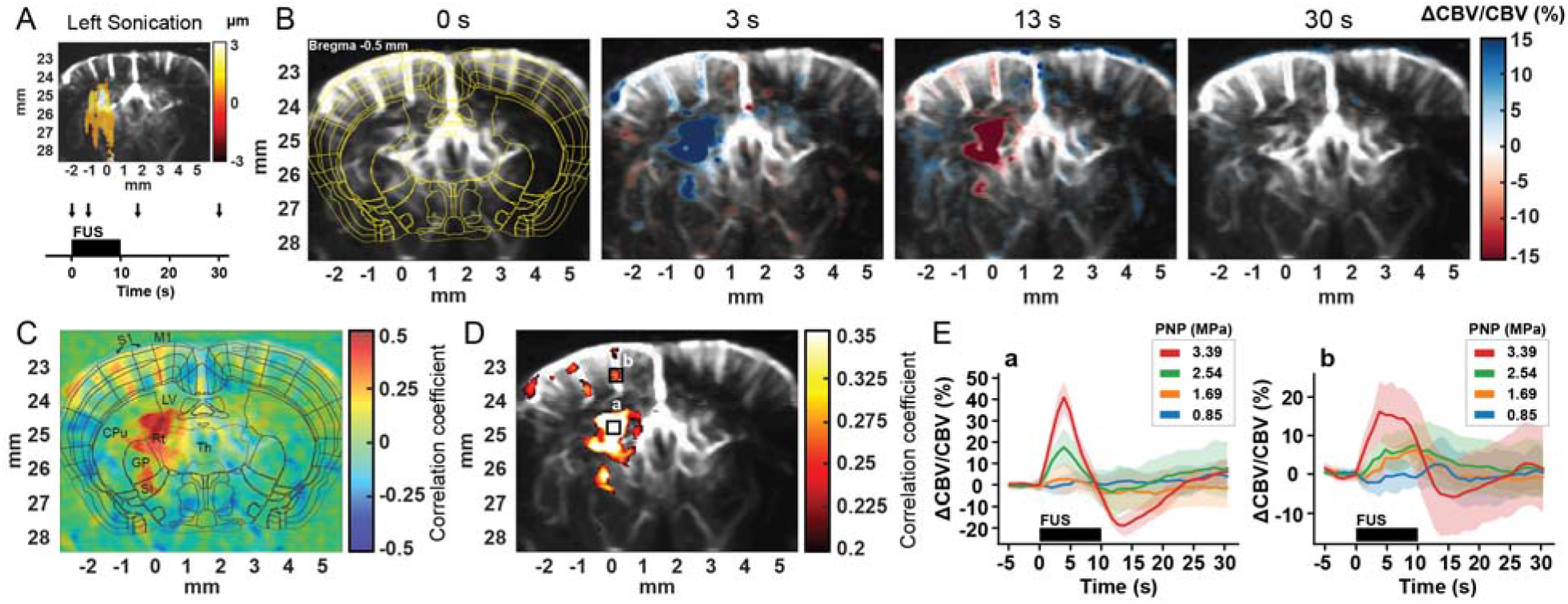
Functional responses to central FUS stimulation. fUS reveals that FUS evokes hemodynamic responses at the cortical and subcortical structures. (A) Displacement imaging for in situ targeting when the left hemisphere was targeted (top), and FUS sonication timeline (bottom). The arrows denote the time points of CBV images demonstrated in (B). (B) CBV responses evoked by FUS (3.39 MPa, 10 s SD, 300 ms PD) at 0, 3, 13, 30 s after the onset of FUS. (C) Activation map based on the pixel-wise computation of Pearson’s correlation between CBV trace and the binary stimuli signal. S1: Primary somatosensory cortex, M1: Primary motor cortex, LV: Lateral ventricle, CPu: Caudate putamen, Rt: Reticular thalamic nucleus, Th: Thalamus, GPe: Globus pallidus, SI: Substantia innominata. (D) Activated pixels overlaid on the power Doppler image. Two regions of interest (ROIs); a and b represent the subcortical and cortical areas, respectively. (E) Mean CBV responses at ROI a (left) and b (right) at different pressures (PNP range: 0.85-3.39 MPa; SD = 10 s, PD = 300 ms).

### 2.9. Statistical analysis

All statistical analyses were conducted using Prism 9 (GraphPad; San Diego, CA, USA). In characterization of CBV responses to varying FUS parameter, one-way ANOVA with Tukey correction (PNP and SD) and two-tailed paired t-test were performed (PD). To examine colocalization between displacement and hemodynamic activation, a two-tailed nonparametric Spearman correlation and linear regression were performed between Pearson’s correlation coefficient and displacement. The displacement between two pixel groups (activated and non-activated pixels) was statistically analyzed by an unpaired t-test with Welch correction. A two-tail nonparametric Spearman correlation was used to establish the relationship between displacement and FUS-evoked hemodynamic responses.

### 2.10. Histological evaluation

30 minutes after the last sonication was performed, a mouse was sacrificed by transcardial perfusion while under deep anesthesia with an isoflurane/oxygen gas mixture delivered through a nose cone. The animal was placed in a supine position, chest fur clipped, and a thoracotomy was performed to expose the chest cavity to access the heart. A needle was placed in the left ventricle and the right atrium was clipped while 1x PBS was flushed at a rate of 0.3 mL/minute for 5 minutes until exsanguinated. Brain tissue was collected by decapitation and placed in 4% paraformaldehyde (PFA) for 48-72 hours, then switched to a series of ethanol incubations for 48 hours and stored in 70% ethanol until the specimen was submitted to a histology core for paraffin-embedding and Hematoxylin and Eosin (H&E) stain. To investigate safety, regions of interest for histological evaluation were chosen as the cortex, caudate putamen, and thalamus

## Results

### 3.1. Displacement imaging can target and monitor tissue displacement in the mouse brain

First, we validated the targeting of 4 MHz FUS using displacement imaging and measured displacement in the brain. Displacement imaging was performed using 3 ms and 2.54 MPa FUS pulse. Fig. 2A depicts a coronal B-mode image of craniotomized mouse brain at Bregma -0.5 mm. The target FUS pulse location (center sonication) matches the high displacing area in the displacement images which is expected because the ultrasound beam energy is the highest at the focus. This demonstrates that displacement imaging can be used to locate the FUS pulse application area. See Fig. 2B, Supplementary Video S3 for detailed of the changes in the displacements during the on-off period of the FUS pulse. Positive and red-colored displacements mean downward or away from the imaging transducer displacement due to the FUS pulse whereas negative and blue-colored means upward or toward the imaging transducer displacements due to the relaxation of brain tissue experiences during FUS off period. Fig. 2C depicts interframe and cumulative displacement traces. The maximum amplitude of interframe and cumulative displacement were found to be 2 µm and 3 µm, respectively.

### 3.2. FUS evokes hemodynamic responses at cortical and subcortical structures

Fig. 3A depicts a displacement map with left sonication and the arrows denote the time points of CBV images depicted in Fig. 3B. Cortical and subcortical regions in the sonicated hemisphere exhibit an increase in CBV followed by its decrease and it recovered to the baseline (Fig. 3B; Supplementary Video S4.1 and S4.2). Note that, we consistently observed this ipsilateral hemodynamic response to FUS in all the mice used in our experiment. To identify brain structures that exhibited CBV increases in response to FUS, we registered stereotaxic brain maps (Allen Brain Atlas; Allen Institute, Seattle, WA, USA) onto functional ultrasound images and activation maps. Activation map reveals CBV increases in primary sensory (S1) and motor (M1) cortex, and subcortical structures including caudate putamen (CPu), globus pallidus (GP), reticular thalamic nucleus (Rt), and thalamus (Fig. 3C). Higher correlation (i.e. stronger activation) was observed in the subcortex compared with the cortex. Fig. 3E depicts mean CBV changes from one animal (n = 1) in two ROIs (Fig. 3D) before, during and after FUS (10 s SD, 300 ms PD, PNP denoted in the figure). At ROI a (subcortex), CBV increases, peaks at 4s with a peak ΔCBV/CBV of 42.6 ± 7.5 % at 3.39 MPa (Mean ± STD), and recovers to the baseline following an undershoot. At ROI b (cortex), CBV increases, peaks at 4s with a lower peak ΔCBV/CBV of 16.4 ± 7.9 % at 3.39 MPa (Mean ± STD) and a plateau, and returns to the baseline. In addition, we observed pressure-dependent hemodynamic activation and higher pressure results in a higher CBV increase. Therefore, we next asked how ultrasonic parameters affect FUS-evoked hemodynamic responses.

### 3.3. CBV responses are dependent on FUS parameters

To characterize dose-dependent hemodynamic response, we explored different combination of ultrasonic parameters including peak negative pressure (PNP), sonication (SD) and pulse duration (PD) over sessions (Supplementary Table S2). Here, we quantitatively evaluated subcortical CBV responses at ROI a. Peak ΔCBV/CBV, activated area, and maximum correlation coefficient from all animals (n = 4) were quantified. We observed monotonically increasing peak ΔCBV/CBV, activated area, and maximum correlation coefficient with increasing PNP (Fig. 4(A1), PNP range: 0.85-3.39 MPa; SD = 10 s, PD = 300 ms). Peak ΔCBV/CBV values were 4.28 ± 1.29 %, 3.14 ± 0.98 %, 10.14 ± 3.02 %, and 25.92 ± 6.07 % with 0.85, 1.69, 2.54, and 3.39 MPa PNP, respectively (Fig. 4(A2), Mean ± SEM, one-way ANOVA with Tukey correction). Activated area values were 0.02 ± 0.01 *mm*^2^, 0.13 ± 0.04 *mm*^2^, 0.31 ± 0.06 *mm*^2^, and 1.5 ± 0.4 *mm*^2^ with 0.85, 1.69, 2.54, and 3.39 MPa PNP, respectively (Fig. 4(A3), Mean ± SEM, one-way ANOVA with Tukey correction). Maximum correlation coefficient values were 0.20 ± 0.02, 0.27 ± 0.02, 0.32 ± 0.02, and 0.40 ± 0.03 for 0.85, 1.69, 2.54, and 3.39 MPa PNP, respectively (Fig. 4(A4), Mean ± SEM, one-way ANOVA with Tukey correction). As shown in Fig. 4(B1), peak ΔCBV/CBV, activated area, and maximum correlation coefficient also monotonically increase with SD (SD range: 1-10 s; PNP = 3.39 MPa, PD = 300 ms). Peak ΔCBV/CBV values were 2.42 ± 0.66 %, 6.36 ± 1.28 %, and 25.50 ± 6.77 % with 1, 5, and 10 s SD, respectively (Fig. 4(B2), Mean ± SEM, one-way ANOVA with Tukey correction). Activated area values were 0 ± 0 *mm*^2^, 0.95 ± 0.81 *mm*^2^, and 2.17 ± 1.34 *mm*^2^ with 1, 5, and 10 s SD, respectively (Fig. 4(B3), Mean ± SEM, one-way ANOVA with Tukey correction). Maximum correlation coefficient values were 0.12 ± 0.01, 0.35 ± 0.07, and 0.45 ± 0.06 with 1, 5, 10 s SD, respectively (Fig. 4(B4), Mean ± SEM, one-way ANOVA with Tukey correction). Fig. 4(C1) shows greater CBV responses at 300 ms PD with significantly higher peak ΔCBV/CBV area and maximum correlation coefficient, compared with 150 ms PD (PNP = 3.39 MPa, SD = 10 s). Peak ΔCBV/CBV values were 13.47 ± 3.3 and 32.79 ± 7.08 % with 150 and 300 ms PD, respectively (Fig. 4(C2), Mean ± SEM, two-tailed paired t-test). Activated area values were 0.74 ± 0.5 *mm*^2^, and 2.47 ± 1.17 *mm*^2^ with 150 and 300 ms PD, respectively, (Fig. 4(C3), Mean ± SEM, n = 4, two-tailed paired t-test). Maximum correlation coefficient values were 0.31 ± 0.03, and 0.46 ± 0.05 with 150 and 300 ms PD, respectively (Fig. 4(C4), Mean ± SEM, two-tailed paired t-test). The results indicate that FUS-evoked hemodynamic responses are dependent on pressure, total sonication duration, and pulse duration. Activation maps according to each parameter set are demonstrated in Supplementary Fig. S1.

**Figure 4.**
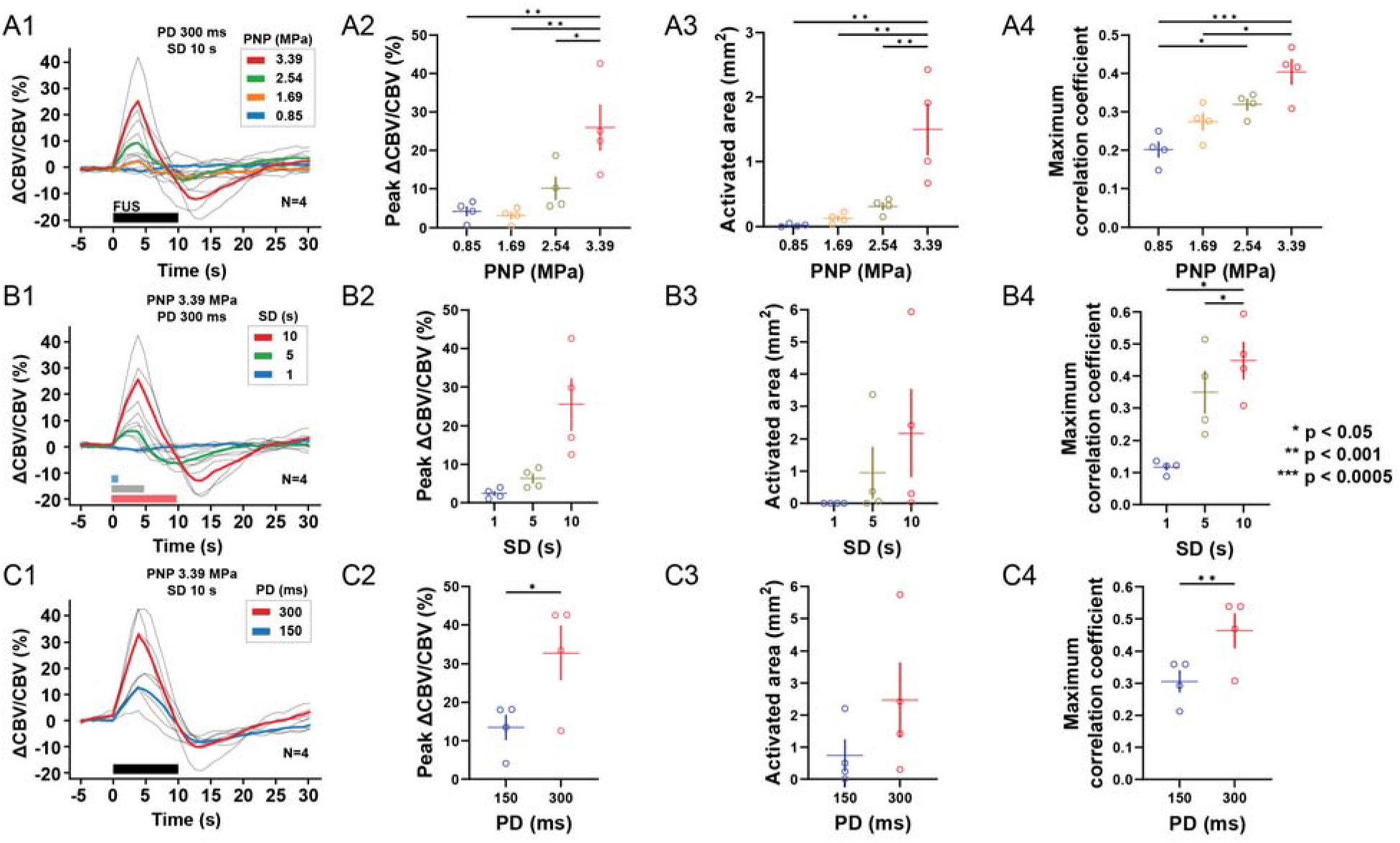
Characterization of CBV responses to varying FUS parameters (PNP, SD, and PD, n = 4 animals). (A1) CBV responses to varying PNP values (SD = 10s, PD = 300ms). Statistical results (Mean ± SEM with data) on (A2) peak CBV changes, (A3) activated area, (A4) maximum correlation coefficient with different PNP values. (B1) CBV responses to varying SD values (PNP =3.39 MPa, PD = 300ms). Statistical results (Mean ± SEM with data) on (B2) peak CBV changes, (B3) activated area, (B4) maximum correlation coefficient with different SD values. (C1) CBV responses to varying PD values (PNP = 3.39 MPa, SD = 10s. Statistical results (Mean ± SEM with data) on (C2) peak CBV changes, (C3) activated area, (C4) maximum correlation coefficient with different PD values. For statistical analysis in Fig. 4A and 4B, one-way ANOVA with Tukey correction was performed for multiple comparisons. For statistical analysis in Fig. 4C, a two-tailed paired t-test was performed. Any significance in the performed analyses is presented (*p < 0.05, **p < 0.001, *** p < 0.0005).

### 3.4. Lateralized target engagements result in lateralized hemodynamic responses

As we leveraged high spatial resolution of 4 MHz FUS, we expected spatially localized hemodynamic activation somewhat reflecting the size of FUS focus or displacement map. To verify this, fUS imaging was conducted with FUS targeted at the three locations (see Materials and Methods). Fig. 5 depicts the activation maps with left, center, and right sonication (column) over increasing pressures (row). The activation map showed lateralized hemodynamic activation in the cortex and subcortex according to the sonicated side. We observed ipsilateral hemodynamic responses with the left and right sonication and bilateral hemodynamic response with the center sonication. The number of activated pixels increases with pressure and the center sonication showed the greatest number of activated pixels.

**Figure 5.**
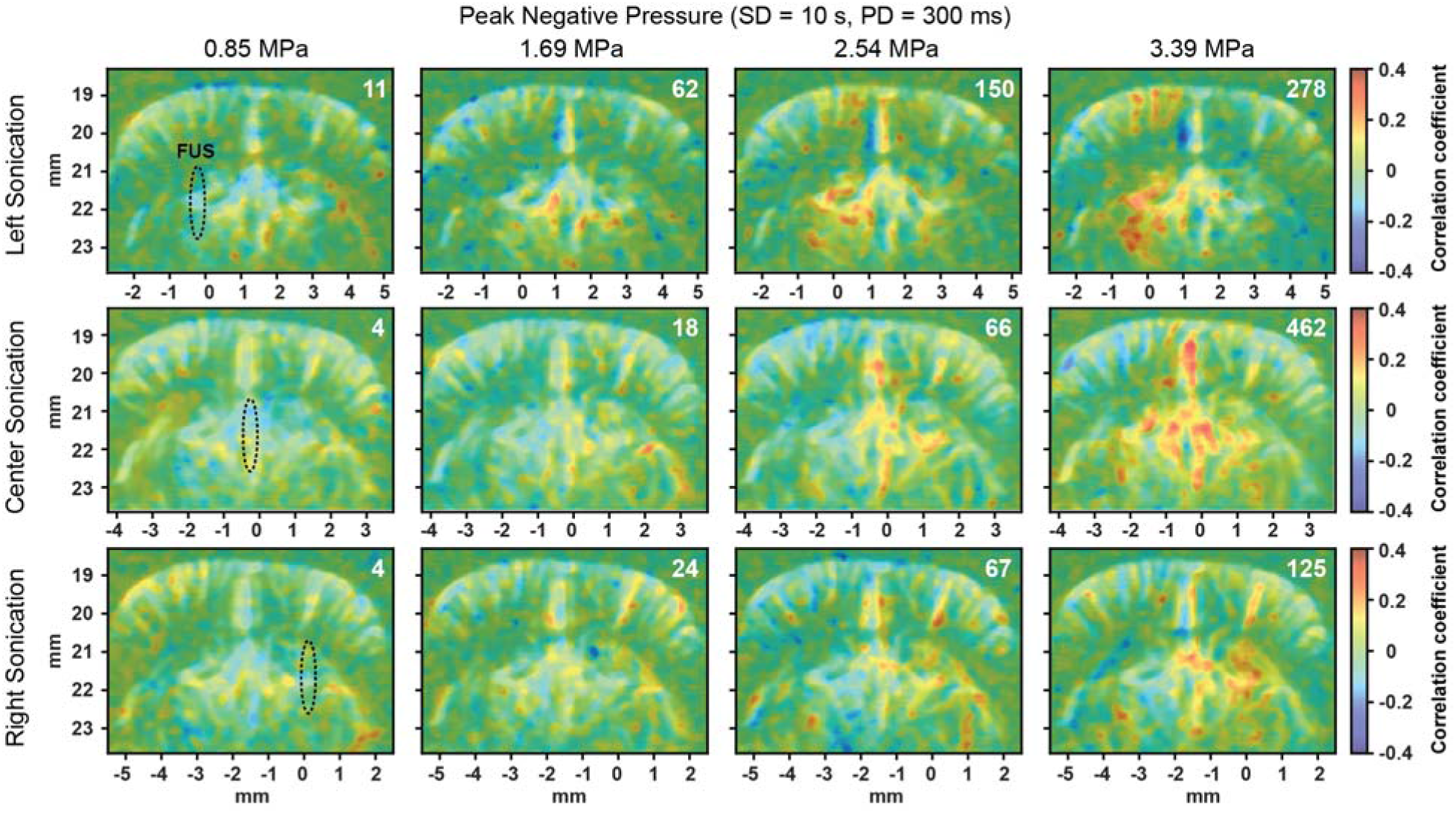
Lateralization of FUS-evoked hemodynamic responses. 12 activation maps from one animal with laterally different target engagements with increasing pressures. The first, second, and third rows depict the activation maps with the left, center, and right sonication, respectively. The dotted area overlaid on the activation map depicts the full width at half maximum (FWHM) of FUS focus. The number on the top right in the activation map denotes the number of activated pixels.

### 3.5. Hemodynamic activation correlates and colocalizes with displacement

Finally, we investigated correlation and colocalization between hemodynamic activation and displacement. Fig. 6A depicts a displacement-activation map with the left sonication (Supplementary Video S5). To quantitatively evaluate the colocalization between hemodynamic response and displacement, we performed a two-tailed nonparametric Spearman correlation between displacement and Pearson’s correlation coefficient of all pixels in the brain (Fig. 6B; 12565 pixels). A pixel with higher displacement showed a greater correlation coefficient (p = 0.0011), suggesting FUS-evoked hemodynamic activation colocalizes with displacement. The group of the activated pixels showed significantly greater displacement than the group of non-activated pixels as shown in Fig. 6C (p < 0.0001, unpaired t-test with Welch correction). The displacement of activated pixels and non-activated pixels were 1.02 ± 0.49 µm, 0.14 ± 0.55 µm, respectively (Mean ± STD). Next, we examined the correlation between peak ΔCBV/CBV and displacement. CBV responses and the FWHM averaged displacement with different pressures (PNP range: 0.85-3.39 MPa; SD = 10 s, PD = 300 ms) were used in the analysis (n = 4). Fig. 6D shows higher displacement results in stronger CBV responses (p = 0.0115, two-tailed Spearman correlation) and CBV responses linearly increase as displacement increases (R = 0.7552, linear regression).

**Figure 6.**
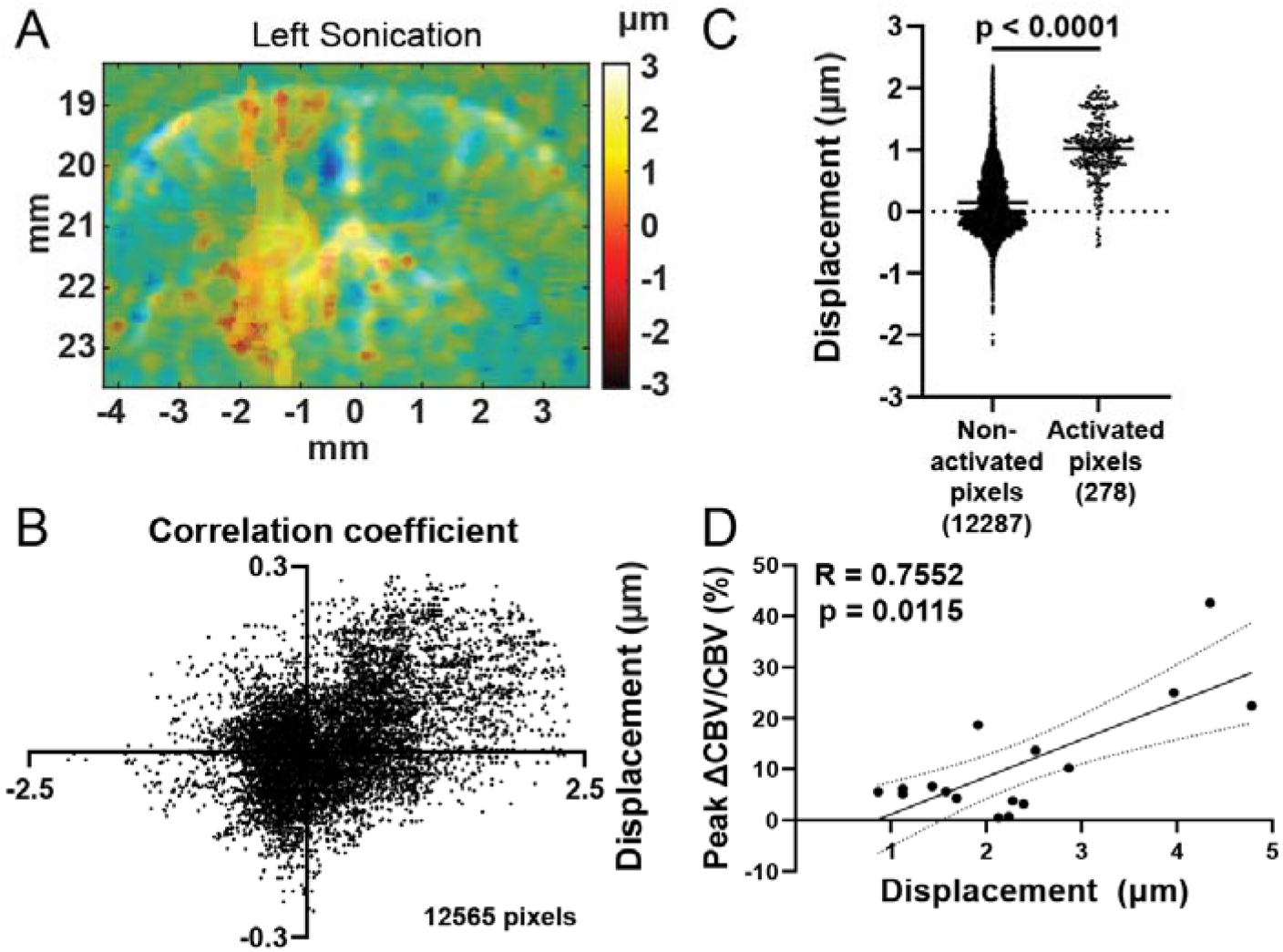
Displacement-activation map reveals colocalization and correlation between displacement and CBV response. (A) Displacement-activation map with left sonication. (B) Scatter plot of correlation coefficient and displacement. Correlation between displacement and correlation coefficient was evaluated by computing a two-tailed nonparametric Spearman correlation (p = 0.0011). (C) Comparison of displacement between two groups of pixels; one consists of 12287 non-activated pixels, and the other consists of 278 activated pixels (n = 1). An unpaired t-test with Welch’s correction was performed to evaluate significance (p < 0.0001). (D) Correlation between Peak CBV increase and displacement. A total of 16 CBV responses with different pressures were used in the analysis (n = 4). Linear regression was performed (R = 0.7552), and a two-tailed nonparametric Spearman correlation was computed (p = 0.0115).

## Discussion

Elucidating how FUS displaces structures in the brain and affects hemodynamics is important to shed light on the mechanism of FUS and also ensure successful targeting and neuromodulation. In this study, we introduced a fully ultrasonic approach to in situ target via displacement, modulate brain activity via FUS, and assess subsequent neuromodulation outcomes via fUS. fUS imaging enables whole-brain neuroimaging of FUS-evoked hemodynamics, allowing spatial resolutions of 100 µm and temporal resolutions of 1 Hz which were enough to find out correlated blood volume and obtain hemodynamic activation maps.

We repeatedly obtained ipsilateral hemodynamic responses in the cortex and subcortex depending on lateralized sonication. The area of the observed responses spans the somatosensory and motor cortex and subcortical regions of caudate putamen, globus pallidus, reticular thalamic nucleus, and thalamus. These subcortical structures are central components of basal ganglia known to govern motor, emotional, and cognitive function [35]. Thus, our findings highlight the importance of neuroimaging of deep brain structures during FUS, which was successfully achieved with fUS imaging in this study. We did not observe any motor responses at the limb and tail as many previous studies reported [6,9,11-14,25,36]. This may be largely due to the difference in the type of anesthetic and the level of anesthesia. Since isoflurane induces vasodilation [38], leading to a lower contrast of fUS signals to the baseline, we herein maintained 0.8 – 1 % isoflurane to keep animals stable during fUS imaging and FUS. Under the modest level of anesthesia, we were able to prevent motion artifacts into power Doppler images, but also acquire fUS signals enough to detect CBV changes evoked by FUS. The reported motor responses were prominent under low-levels of isoflurane (< 0.1 %) close to awake state [11,36], sometimes with introducing wash-out period to allow the animal to recover from anesthesia [37]. Given that the type of anesthetic affects hemodynamic activation pattern [39], it would be beneficial to investigate the effect of different anesthetics and doses on FUS-evoked hemodynamic response.

fUS imaging reveals distinct responses to FUS between cortical and subcortical regions with different hemodynamic functions. It may be due to the higher intensity of ultrasound at a subcortical area close to the depth of our target location (DV: 3 mm), but we also hypothesize that subcortical region is more susceptible to FUS-mediated neuromodulation due to its lower stiffness than cortical region. To investigate brain stiffness maps, we performed single-transducer harmonic motion imaging (ST-HMI) on a craniotomized mouse [40]. Interestingly, our preliminary data on brain stiffness reveals subcortical structures have larger peak-to-peak displacement induced by FUS, indicating lower stiffness compared with cortical structures (Supplementary Fig S2). This finding is consistent with our observation of a stronger FUS-evoked hemodynamic response at the subcortex compared with the cortex. Given that FUS-evoked neuronal activation is mediated by mechanosensitive channels triggered by acoustic radiation force (or displacement), we hypothesize the subcortex is highly susceptible to FUS because the subcortical area can be displaced larger than the cortical area at a given pressure. It demonstrates the importance of displacement imaging in the context of mechanistic monitoring of FUS.

We recently reported hemodynamic responses evoked by FUS in mice and nonhuman primates via transcranial fUS [41]. The study reported bilateral responses with center sonication and less localized hemodynamic responses with left or right sonication, which may be largely due to a large focal size of 1.68 MHz FUS transducer without in situ displacement targeting, and the presence of the skull. Compared with the study, we herein were able to achieve more targeted FUS neuromodulation by utilizing high frequency FUS and displacement imaging, and eventually reveal colocalization and correlation between CBV and displacement.

How do we interpret hemodynamic responses observed in this study and compare them with the results reported in previous studies? We saw both the cortical and subcortical CBV peak at 4 s which is comparable to or slower than 3.2 s in rabbits [7], 2.5 s in mice [26], and 2.7 s in mice [27]. Interestingly, our fUS imaging reveals undershoot (negative CBV), followed by CBV returning to baseline, which none of the aforementioned studies reported, but BOLD fMRI studies reported as negative BOLD or post-stimulus undershoot [42-44]. Our study and [27] showed CBV peaks and begins to decline during FUS while [7] and [26] showed CBV peaks post-FUS. However, it may be difficult to directly compare those results because of the difference in sonication parameters and brain imaging modalities (fMRI BOLD [7], optics [26,27], fUS: this work). The difference in FUS-evoked hemodynamic responses may arise due to the difference in sensitivity in imaging modalities, ultrasonic parameters, and anesthesia conditions.

We found that the FUS-mediated CBV increase is dependent on ultrasonic parameters including pressure, total sonication duration and pulse duration (Fig. 4). The range of ultrasonic parameter explored in this study was chosen based on FUS stimulation paradigms used in our previous work on central FUS stimulation in mice [13,14]. Interestingly, peak ΔCBV/CBV and activated area appears to non-linearly increase with pressure. This observation is consistent with the acoustic radiation force-based mechanism of FUS stimulation, given that the acoustic radiation force is proportional to approximately the square of the pressure. Our finding on dose-dependency of hemodynamic activation induced by FUS is in good agreement with [26], where the researchers showed FUS-evoked hemodynamic response is dependent on pressures and duration in mice. We did not observe robust CBV increases at short sonication duration of 1s, however they did observe CBV response at even lower sonication duration and intensity (SD = 400 ms, *f_SPPA_* = 1.1 W*/cm*^2^). It might be due to the relatively lower fUS sensitivity with the imaging transducer (f=15.625 MHz) used in this study. It may be addressed by using a high-frequency imaging transducer (f=40 MHz) capable of capturing smaller vessels with a diameter up to 75 µm [24], but it was not utilized in this study due to incompatible geometry of the FUS and high-frequency imaging transducers. It is intriguing to see lateralized and dose-dependent hemodynamic activation via correlation map depicted in Fig 5 and Supplementary Fig. S1. Overall, the size of the activated area and the correlation coefficient increases as ultrasonic dose increases. The number of the activated pixels was greatest in case of center sonication. This may be because the center sonication may cause bilateral brain activation previously reported as bilateral cerebral blood flow (CBF) increases [27], bilateral limbic [14], and bilateral whisker movement [26]. Interestingly, our data may indicate that the activated area is not necessarily confined to the focal area, usually larger than the focal area, which may be plausible given that entire brain tissue surrounding the focus can be displaced by FUS along the beam path. It underlines the merits of imaging displacement and functional response of the entire brain to assess the neuromodulatory effect of FUS.

In this study, we adopted an amplitude-modulated pulsing scheme at 1 kHz AM frequency to minimize auditory confounding effect that could occur during sonication [3]. Recently, non-the auditory confounding effect has been also reported [45], which may be caused by non-auditory stimuli (i.e. tactile and visual stimuli) evoked by FUS. However, our finding of dose-dependent and lateralized hemodynamic activation in response to FUS discounts that the hemodynamic response is attributed to the confounds. Thus, we ascertain that the CBV responses we observed and reported in this study were driven by FUS, not any confounding effects.

To assess the safety of FUS protocol used in this study, we sonicated the left hemisphere of a craniotomized mouse with consecutive 3 sessions of fUS-FUS with the parameter set (a total of 21 sonications with PNP = 3.39 MPa, SD = 10 s, PD = 300 ms) that can induce maximum ultrasonic dose among the parameter sets we used in the study and performed H&E staining. We did not detect any direct damage or red blood cell extravasation in both hemispheres (Supplementary Fig S3).

This study has the limitation that we cannot completely separate the thermal effect of ultrasound on hemodynamic activation from the mechanical effect. Thermocouple-measured temperature rise at the subcortical region was reported in Supplementary Table S3. The maximum temperature rise was 5.24 °C at the end of sonication (PNP = 3.39 MPa, SD = 10 s, PD = 300 ms). Temperature rise can excite cells [46] and the mechanosensitive Piezo2 channel was reported less active at lower temperatures [47]. Under the ultrasonic parameters we used, both thermal and mechanical effects could plausibly contribute to the neuromodulatory effect of ultrasound, which can be accepted as long as the thermal dose does not exceed the threshold of tissue damage. As stated previously, we have confirmed none of the sonication parameters or cumulative effects that could occur during the experiment did not induce any significant damage. Our previous work [13] demonstrating ipsilateral and contralateral limb movement selectively may be an exemplar of temperature-facilitated FUS neuromodulation. Under the FUS parameter used in the study, temperature elevation was found to be 6.8 °C [14]. We hypothesize that the highly selective motor responses may be able to be achieved with temperature rise. In temperature measurement, we observed monotonically increasing temperature during sonication. A hemodynamic pattern we observed that CBV peaks and starts decreasing in the middle of sonication may suggest the hemodynamic response is not likely mainly driven by the thermal effect. Yet, still some of the hemodynamic responses might arise from the temperature elevation.

Displacement imaging provides a displacement map with spatial resolutions of 100 µm and temporal resolutions of 0.72 ms. We were able to successfully image the acoustic radiation force induced by FUS and demonstrated the feasibility of in situ confirmation of FUS targeting via displacement. Brain tissue displacement at 2.54 MPa was ∼ 2 µm, which is a comparable amplitude of displacement measured in a large animal model with MR-ARFI [15]. By mapping displacement and activation maps, we found that displacement colocalizes and linearly correlates with CBV increase. This finding is consistent with earlier two studies on the relationship between the neuromodulatory impact of FUS and displacement [5,15]. Menz et al. [5] reported that neural spiking activities of *ex vivo* retina correlate with optically imaged displacement and Mohammadjavadi et al. [15] found a suppressive effect of FUS on visual-evoked potential correlates with MR-ARFI displacement. Furthermore, our displacement-activation map may indicate the tissue displacement required for neuronal activation is above 1 µm (Fig. 6C). The findings presented herein on the relationship between displacement and CBV provide another *in vivo* evidence of acoustic radiation force-based mechanism of FUS neuromodulation.

While our study probes the hemodynamic activation map in response to FUS, we only recorded acute CBV changes in the fixed imaging plane. A longitudinal study would be compelling to investigate the offline effect of FUS and functional connectivity changes [48]. Whole-brain neuroimaging [49] will provide volumetric CBV allowing us to investigate ascending and descending pathways or brain networks possibly triggered by FUS [19,22]. Therefore, we can consolidate our understanding of the neuromodulatory effect of FUS in the brain.

## Conclusion

This study first introduced a fully ultrasonic approach to in situ target via displacement imaging, modulate neuronal activity via FUS, and monitor the resultant neuromodulation effect via fUS imaging. We demonstrated displacement imaging on craniotomized mice and its feasibility as a way of in situ confirmation for targeting. We saw ipsilateral hemodynamic increases in CBV that peak at 4 s. Importantly, we saw a stronger correlation at the subcortical area than cortical area. The finding is consistent with our preliminary data on the brain elasticity map and we hypothesize that the subcortical region is highly susceptible to FUS-mediated neuromodulation due to its lower stiffness. We found that CBV response is dependent on pressure, total sonication duration and pulse duration and the highest CBV increase was 25.92 ± 6.07 % (Mean ± SEM; PNP = 3.39 MPa, SD = 10 s, PD = 300 ms). Peak ΔCBV/CBV, the size of the activated area and the correlation coefficient increase with ultrasonic dose. The thermocouple-measured temperature was not negligible, but we have confirmed the safety of our FUS protocol used in this study with H&E staining. Additionally, by mapping displacement and hemodynamic activation, we discovered that displacement colocalizes and linearly correlates with CBV. Our data may suggest that the FUS-induced displacement required in evoking neuronal activation is above 1 µm. The findings presented in this study will help guide future studies of hemodynamic responses to FUS and by elucidating displacement as a result of the acoustic radiation force by which FUS exerts on neuronal tissue, we anticipate that the insights developed in this study will help inform more guided FUS neuromodulation.

## Acknowledgement

This work was funded by National Institute of Health (RO1EB027576). The authors would like to thank Pablo Abreu, M.A., for his gracious assistance, financially and administratively; Stephen A. Lee, Ph.D. and Christian Aurup, Ph.D., for their mentorship and training of FUS and fUS imaging; Erica P. McCune, M.S., Samuel G. Blackman, B.S, Aparna Singh, Ph.D., Sergio Jiménez-Gambín, Ph.D. for their insightful discussion.

