## Supplementary Material for "Functional ultrasound (fUS) imaging of displacement-guided focused ultrasound (FUS) neuromodulation in mice"

Supplementary Table S1. Imaging parameters for displacement and functional ultrasound imaging.

| Category | Parameter | Value |
| --- | --- | --- |
| Displacement imaging | Imaging array (frequency) | L22-14vX-LF (15.625 MHz) |
|  | Number of angles | 12 |
|  | Maximum angle | ± 3° |
|  | Framerate | 16667 Hz |
|  | FUS pulse duration | 3 ms |
| fUS imaging | Imaging array (frequency) | L22-14vX-LF (15.625 MHz) |
|  | Number of angles | 13 |
|  | Maximum angle | ± 7° |
|  | Number of RF signals averaged per angle | 3 |
|  | Number of compound frames | 150 |
|  | Framerate | 19500 Hz |
|  | SVD cutoff | 30 |
|  | Power Doppler framerate | 1 Hz |

Supplementary Table S2. Combination of FUS parameters used in this study.

| Session # | PNP (MPa) | SD (s) | PD (ms) | $I_{SPPA} (W/{cm}^{2})$ | MI ($PNP/\sqrt{f}$) |
| --- | --- | --- | --- | --- | --- |
| 1 | 0.85 | 1 | 300 | 23 | 0.42 |
| 2 | 1.69 | 1 | 300 | 89 | 0.85 |
| 3 | 2.54 | 1 | 300 | 202 | 1.27 |
| 4 | 3.39 | 1 | 300 | 360 | 1.69 |
| 5 | 0.85 | 5 | 300 | 23 | 0.42 |
| 6 | 1.69 | 5 | 300 | 89 | 0.85 |
| 7 | 2.54 | 5 | 300 | 202 | 1.27 |
| 8 | 3.39 | 5 | 300 | 360 | 1.69 |
| 9 | 0.85 | 10 | 300 | 23 | 0.42 |
| 10 | 1.69 | 10 | 300 | 89 | 0.85 |
| 11 | 2.54 | 10 | 300 | 202 | 1.27 |
| 12 | 3.39 | 10 | 300 | 360 | 1.69 |
| 13 | 3.39 | 10 | 150 | 360 | 1.69 |

Supplementary Table S3. Peak temperature increases under parameter sets used in this study.

| Temperature rise | | PNP (MPa) | | | |
| --- | --- | --- | --- | --- | --- |
| PD (ms) | SD (s) | 0.85 | 1.69 | 2.54 | 3.39 |
| 150 | 10 | N/A | | | 2.78 °C |
| 300 | 1 |  |  |  | 1.56 °C |
| 300 | 5 | 0.31 °C | 1.05 °C | 2.02 °C | 3.74 °C |
| 300 | 10 | 0.62 °C | 1.72 °C | 3.4 °C | 5.24 °C |

Supplementary Fig S1. Activation maps according to FUS parameters with left sonication. The parameter used was displayed on top of the map (PNP / SD / PD).

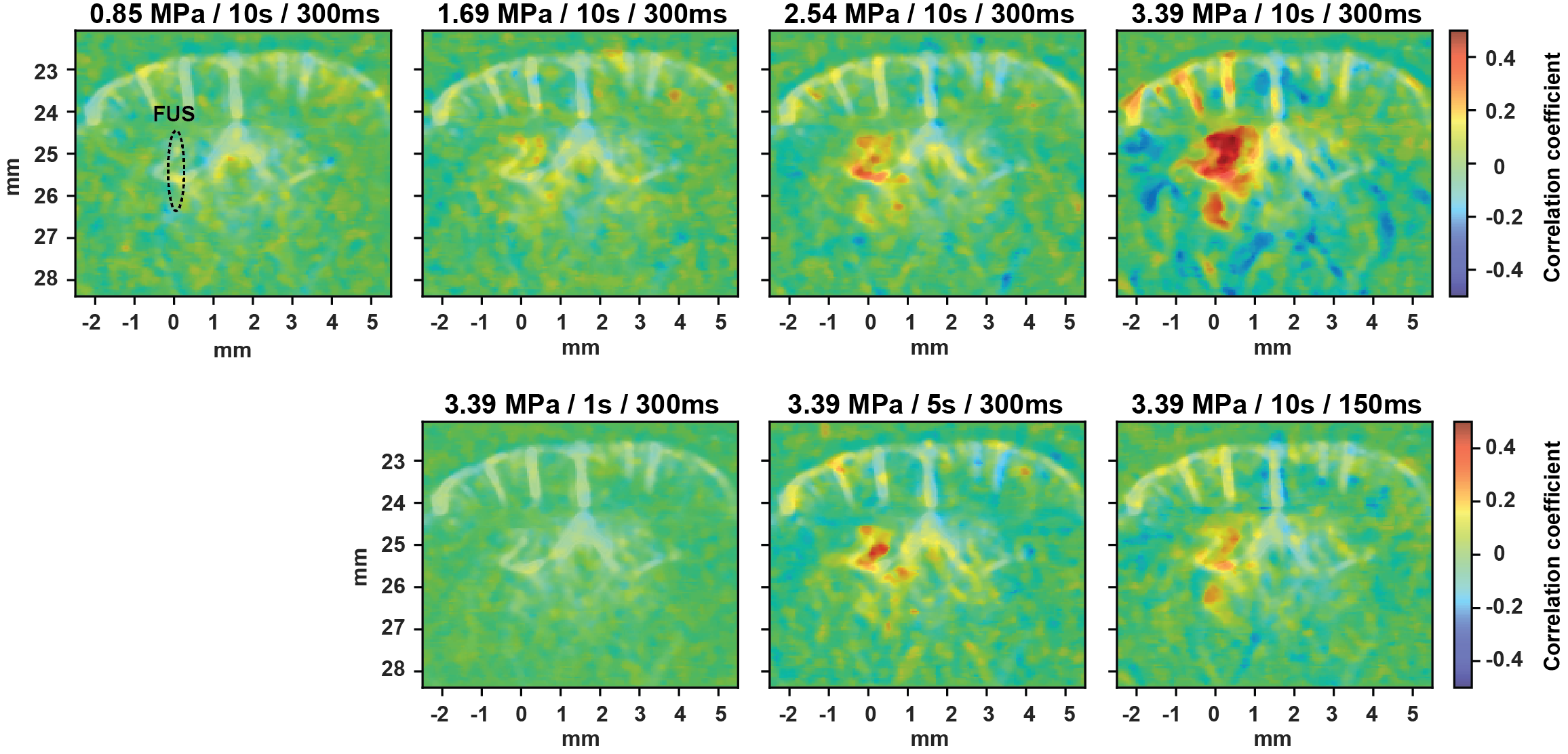

Supplementary Fig S2. Brain elasticity map using ST-HMI. ST-HMI reveals a higher peak to peak displacement ratio at subcortical region in a craniotomized mouse, indicating the subcortical region is softer (i.e. has lower Young’s modulus) than the cortical.

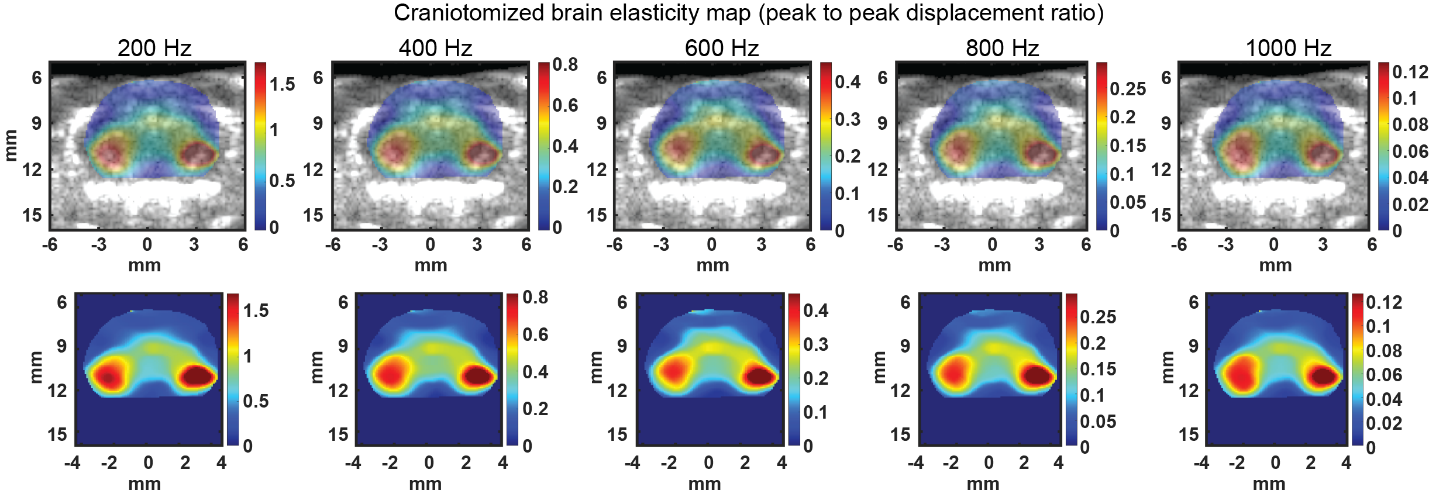

Supplementary Fig S3. Histological analysis (H&E staining). Left hemisphere was sonicated and no significant tissue damages was observed in any of hemispheres.

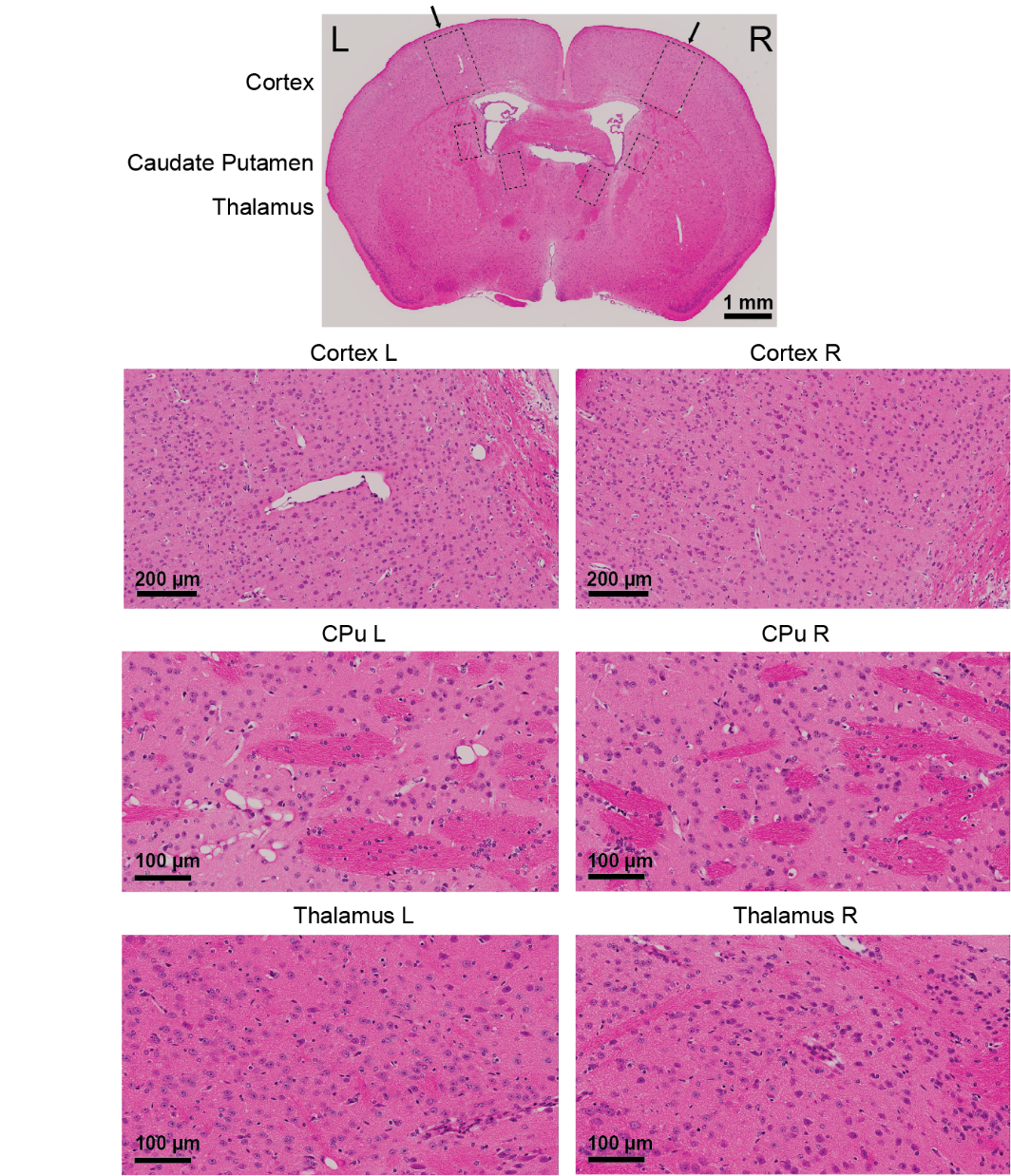
